# Single-cell transcriptomic analysis of immune cell dynamics in the healthy human endometrium

**DOI:** 10.1101/2023.11.09.566466

**Authors:** Kaixing Chen, Qiaoni Yu, Qing Sha, Junyu Wang, Jingwen Fang, Xin Li, Xiaokun Shen, Binqing Fu, Chuang Guo

**Affiliations:** Department of Rheumatology and Immunology, the First Affiliated Hospital of USTC, Division of Life Sciences and Medicine, University of Science and Technology of China, Hefei, Anhui 230021, China; CAS Center for Excellence in Molecular Cell Sciences, the CAS Key Laboratory of Innate Immunity and Chronic Disease, University of Science and Technology of China, 230027 Hefei, Anhui, China; HanGene Biotech, Xiaoshan Innovation Polis, Hangzhou, Zhejiang 311200, China; Department of Rheumatology, The First Affiliated Hospital of Jinzhou Medical University, Jinzhou 121001, China; Department of Immunology, School of Basic Medical Science, Jinzhou Medical University, Jinzhou 121001, China

**Keywords:** Endometrium, Reproductive cycle, Single-cell RNA sequencing, Immune cells, Cell-cell interactions, Reproductive diseases

## Abstract

The microenvironment of the endometrial immune system is crucial to the success of placental implantation and healthy pregnancy. However, the functionalities of immune cells across various stages of the reproductive cycle have yet to be fully comprehended. To address this, we conducted advanced bioinformatic analyses on 230,049 high-quality single-cell transcriptomes from healthy endometrial samples obtained during the proliferative, secretory, early pregnancy, and late pregnancy stages. Our investigation revealed that proliferative natural killer (NK) cells, a potential source of endometrial NK cells, exhibit the most robust proliferative and differentiation potential during non-pregnant stages. During early pregnancy, NK cells display high oxidative phosphorylation metabolism activity, and together with macrophages and T cells, exhibit a strong type II interferon response. Based on our cell-cell interaction analyses, we identify a large majority of interaction pairs to occur in late pregnancy. Finally, we explored the correlation between stage-specific alterations in transcriptomics and the risk genes of common reproductive diseases, unveiling that MHC class I/II molecules, along with *TGFBR1*, exhibited the potential to serve as biomarkers. Our study provides insights into the dynamics of the endometrial immune microenvironment during different reproductive cycle stages, thus serving as a reference for detecting pathological changes during pregnancy.

## Introduction

The reproductive cycle can be broadly classified into the menstrual cycle, which encompasses the proliferative and secretory stages^1^, and the stages of pregnancy, including the early, mid-, and late pregnancy^2^. Many significant occurrences in the reproductive process are closely linked with the endometrium, including embryo implantation, pregnancy, and labor. Such instances are accompanied by a dynamic process involving shedding, regeneration, and differentiation of the endometrial tissue^3^. A comprehensive characterization of the endometrium in healthy individuals throughout the reproductive cycle can facilitate the understanding of the normal transitions and variations in the endometrial microenvironment during different stages of the reproductive cycle, and serve as a foundation for exploring pathological changes in reproductive processes.

The success of pregnancy is greatly influenced by the immune microenvironment of the endometrium^4,5^. As evidenced in a prior study, there is an increase in the proportion of endometrial immune cells from 8.2% during the proliferative stage to 31.7% during early pregnancy^6^. The endometrium contains various types of immune cells, among which the most predominant are NK cells, macrophages, and T cells^7^. During early pregnancy, NK cells comprise 50-70% of leukocytes in normal endometrium^3^. These cells, which could produce and secrete angiogenic factors^8^, play an essential role in instigating and perpetuating decidualization^9^ and facilitating embryo implantation^8,10^. Endometrial macrophages play a vital role in tissue reconstruction, helping to establish a pro-inflammatory microenvironment for embryo implantation, and acting as antigen-presenting cells (APCs)^7^. CD8^+^ cytotoxic T cells can recognize fetal antigens directly through EVTs and indirectly through APCs but would not attack EVTs^11^.

Single-cell RNA sequencing (scRNA-seq) is increasingly used to characterize the endometrial microenvironment at various stages of the reproductive cycle^12^, including identifying cell-cell interactions and new subsets of immune and nonimmune cells^1,13-16^. During early pregnancy, decidual NK cells were observed to comprise three distinct subsets (dNK1, dNK2, and dNK3)^13^. dNK1 are noted for expressing high levels of killer immunoglobulin-like receptors (KIRs) capable of binding to HLA-C molecules. Meanwhile, both dNK1 and dNK2 show the presence of *LILRB1*, which binds to HLA-G molecules, as well as *NKG2A*, which binds to HLA-E molecules. Consequently, dNK1 and dNK2 are believed to potentially interact with EVTs, with dNK1 playing a paramount role. Furthermore, dNK3 and dNK2 are observed to produce more cytokines than dNK1, which implies that they may be more important in recruiting other immune cells and regulating the immune microenvironment^17^. Additionally, recent studies have explored the disparities in the endometrial immune microenvironment between healthy women and those with recurrent pregnancy loss (RPL) during early pregnancy^5,18,19^. However, a comprehensive understanding of the dynamic changes in the endometrial immune environment spanning the reproductive cycle is still lacking.

In this study, we conducted an advanced bioinformatic analysis of large-scale public scRNA-seq datasets obtained from endometrial cells of healthy women across four stages of the reproductive cycle. We identified the stage-specific transcriptomic characteristics of major endometrial immune cells (including NK cells, macrophages, and T cells) during different stages. We also carried out an in-depth analysis of cell-cell interactions and found that there is a strong communication between immune and non-immune cells during different stages of the reproductive cycle, particularly in late pregnancy. Finally, we demonstrated an association between stage-specific transcriptomic changes of endometrial cells and the risk genes of common reproductive diseases. This study provides a relatively comprehensive perspective of the dynamic landscape in the endometrial immune microenvironment throughout the reproductive cycle, which can facilitate studies investigating pathological changes to improve the diagnosis and treatment of endometrium-related ailments.

## Materials and Methods

### Sample information

For the scRNA-seq data we collected, 4 stages of the reproductive cycle were examined: proliferative (n = 4), secretory (n = 11), early pregnancy (n = 28), and late pregnancy (n = 9). For sample sources, 1 of proliferative donors and 1 of secretory donors are deceased organ donors (within 1 h of circulatory arrest) from Garcia-Alonso, L., et al. (*Nat. Genet.*, 2021), and the other 13 of non-pregnant donors are live donors; Decidual tissue of Vento-Tormo et al. (*Nature*, 2018) was obtained from elective terminations of normal pregnancies between 6 and 12 weeks gestation. In the other 3 datasets of early pregnancy, Guo et al. (*Cell Discov.*, 2021), Wang et al. (*Genomics Proteomics Bioinformatics*, 2021), and Chen et al. (*Front. Immunol.*, 2021), decidual tissues were obtained from elective terminations of apparently normal pregnancies and CD45^+^ cells were sorted for further sequencing analysis. Tissues of late pregnancy were obtained immediately after delivery. scRNA-seq data used in our study was obtained with 10x Genomics libraries.

### Single-cell RNA-seq quality control

For the raw sequencing SRA files provided by the articles, fastq-dump (v2.8.0) was used here to download and convert to FASTQ.GZ files, and then Cell Ranger (v4.0.0) and the GRCh38 human reference genome provided by it were used for sequence comparison. For data quality control, in this study, six scRNA-seq datasets were first merged and retained 36,601 standard genes using the Merge function of Seurat^20^ (version 4.0.5), and the data were subsequently partitioned into smaller datasets using the difference in the samples to which they belonged. After preprocessing with Seurat’s NormalizeData and ScaleData functions, this study used DoubletFinder^21^ (version 2.0.3) to screen out double cells with a default setting of 7.5%. Subsequently, cells with detected gene counts between 500 and 6000 and with less than 25% mitochondrial gene expression were retained in this study. In addition, we retained genes expressed in at least 10 cells and simultaneously removed mitochondrial genes. Due to severe batch effects in direct follow-up analysis, different integration approaches were tried in this study, including Seurat, Harmony^22^ (version 0.1.0) and scVI^23^ (version 0.13.0). We finally chose Seurat’s integration results, and the integration process included splitting the dataset (SplitObject), normalizing the data (NormalizeData), obtaining highly expressed genes (FindVariableFeatures), obtaining highly expressed genes with a high number of replicates based on the splitting results (SelectIntegrationFeatures), determining integration anchors (FindIntegrationAnchors), and integrating (IntegrateData), while adjusting the split.by parameter of the SplitObject() function in the Seurat standard integration process, the nfeatures parameter in the FindVariableFeatures and SelectIntegrationFeatures functions, the anchor.features parameter in the FindIntegrationAnchors function, and the IntegrateData function’s sample.tree parameter in the FindIntegrationAnchors function and sample.tree parameter in the IntegrateData function to further optimize the integration results. The integration also included Seurat’s standard process for scRNA-seq downstream data processing and analysis, and this study used the integrated data matrix for normalization (ScaleData), PCA principal component analysis (RunPCA), UMAP downscaling (RunUMAP), calculation of neighborhoods (FindNeighbors), and cell clustering (FindClusters, resolution=1.0).

### Single-cell RNA-seq annotation

In this study, the cell types were broadly classified mainly based on the cell type characteristic genes in the Extended Data provided in the article^13^. Additionally, using the Jaccard index (i.e., the ratio of the intersection size of two sets to the size of the concatenated set), our study calculates the similarity between the natural clusters and the original annotations of the datasets to confirm the cluster-cell type correspondence and further selects the cell clusters with more ambiguous cluster boundaries. For cell clusters that require finer distinction or are difficult to distinguish, we adopt the strategy of removing and then downscaling the cluster annotation, and if there is a serious batch effect in the direct downscaling of clusters, then we use Seurat to integrate before downscaling and then use the original data information and cell type signature genes of the datasets for annotation. If it was difficult to confirm the cell type of the cluster with existing annotations or signature genes, we used the FindMarkers function of Seurat to obtain the differentially expressed genes of the cluster relative to other clusters, identified the cell type of the cluster by reviewing the literature and databases, and finally matched the cell type annotations back to the original dataset using the cell ID. After cell annotation was completed, we removed some cells that included (a) NK cells with high expression of heat shock protein (HSP) genes, which may be due to experimental manipulation causing cellular stress; (b) clusters of cells expressing both T cells and Mac signature genes; and (c) clusters of T cells with high expression of antibody-related proteins.

### Gene expression programs analyses

In our study, NK cells were extracted from the data using annotation information, and NK cells were divided into small datasets of different cell subsets using annotation information in the same way. Then, differentially expressed genes of different stages were obtained for the split small datasets after removing HSP genes (FindAllMarkers, min.pct=0.25, only.pos=TRUE). To explore whether there are some differences between different cell types, this study also used stage information to divide NK cells into small datasets of different stages to obtain differentially expressed genes of different cell subsets. Finally, we merged these genes into one differential gene set. Since some cell subsets of macrophages and T cells were too few in number in some stages, we were concerned that their differential genes would bring huge errors to the whole, so we set at least 30 cells of that cell type in each stage to be retained and then used the retained cell subsets for the above operation, and finally, we obtained three differential gene sets. cNMF can infer gene expression programs, including cell type gene modules and functional gene modules, from scRNA-seq^24^. cNMF will be used in this study to further uncover stage-specific functional transcriptional modules. Since cNMF (version 1.3.2) runs on a python (v3.7.0)-based environment, which requires a matrix file with a.txt format or an object file with a.h5ad format as input, we extracted NK cells from the original Seurat objects after keeping only the genes in the differential gene set. Then, we used the SeuratDisk (version 0.0.0.9019) functions SaveH5Seurat and Convert functions to convert the object and store it as an.h5ad file. The object then undergoes the standard processing flow of cNMF. The main cNMF programs are run in the cnmf_env virtual environment from the command line, which can also be called from the command line in python for ease of subsequent processing. The process includes normalizing the input matrix and preparing the run parameters (prepare instruction), decomposing the matrix (factorize instruction), merging the result file (combine instruction), traversing the result and plotting the stability and error (k_selection_plot instruction), and selecting the result according to the desired k value after presetting the outlier threshold (consensus The -k parameter in the factorize command corresponds to the number of modules of the subsequent genes, and in combination with the existing number of differential genes, we calculate 10, 15 and 20 in NK cells, 8-12 in macrophages and 10-15 in T cells, and use the parallel command to shorten the running time during the calculation. local maximum stability and minimum error is chosen, but there is no clear standard, and the developers suggest using it according to the actual situation. Ultimately, this study obtained the top 50 (NK cells) or top 20 genes (macrophages and T cells) from each gene module based on the results of the gene expression program using the z score.

### Stage-specific gene modules analysis

We use NK cells as an example. In this study, we used the gene modules obtained from the above steps to calculate the average expression of each gene module in each NK cell using the expression matrix after standard processing. Then, using this gene module average expression matrix, we calculated the Pearson correlation coefficient (PCC) (cor, method=Pearson) between the average expression of the 20 gene modules. We then used pheatmap (version 1.0.12, clustering_method=ward. D) to plot the heatmap and obtained 10 gene modules (GMs) based on the hierarchical clustering it provided, and macrophage and T-cell correlations were processed in the same way. To obtain stage-specific or cell type-specific functional transcriptional modules, the acquired GMs were calculated for the expression of cellular isoforms in all stages, and box-line plots were drawn in this study (ggplot2, version 3.3.5). We considered a GM to be stage-specific if it had significantly higher expression (unpaired t test and fold change ≥ 1.1) in at least two cell types at the same identical stage and no other significantly higher-expressing GM occurred in all cell types. We also identified cell type-specific GMs, which may have higher expression in at least three stages than all other cell types. After deduplication of the gene sets in GM, we used clusterProfiler^25^ (version 3.14.3) to perform functional enrichment analysis, with a process including gene ID conversion (bitr, org.Hs.eg.db version 3.10.0) and functional enrichment (enrichGO, ont=ALL). Since only a very small number of genes could not be converted from SYMBOL to ENTREZID during the conversion process, we did not perform subsequent gene ID conversions, and since most of the observed functions were found in BP (biological process), subsequent function enrichment was performed directly using enrichGO (ont=BP, keyType=SYMBOL), which will also facilitate the subsequent viewing of the gene corresponding to a particular function. We then combined the results of all GM function enrichments and selected the top 10 functions of each GM enrichment based on the p value as the main functions to be enriched.

### Cytokine signaling activity prediction

In our study, CytoSig^26^ software was used for the prediction of multiple cytokine signaling activities. The single-cell raw matrices of NK cells, macrophages and T cells were first normalized by the required log2(TPM/10 + 1), and then the processed matrices were output as.txt files, which were converted to.txt.gz files using the gzip command at the command line. In this study, we used.txt.gz files as the standard inputs to perform the signal activity prediction of 51 cytokines in the CytoSig database in the command line, and finally, we selected the .Zscore (regression coefficient/standard error) files as the result for subsequent usage. Then, we selected the cytokines from it for subsequent display. Finally, we likewise plotted boxplots of cytokine signaling activity results using cell subset and stage information.

### Cell-cell interaction prediction

In our study, Nichenet^27^ (Nichenetr, version 1.0.0) was used for cell-to-cell interaction prediction. The ligand-target a priori model, the recipient ligand network and the weighted integration network provided by Nichenet need to be loaded before performing the prediction of intercellular interactions. We obtained Seurat objects containing only the desired cell types, which were later subjected to formal Nichenet analysis. We needed to first define the signal-sending cell types and the signal-receiving cell types and then define the genes in the signal-receiving cell population that might be affected by the ligands expressed by the interacting cells. Here, we used differentially expressed genes in different stages. We then defined the potential ligands in the signal-sending cell types, calculated the ligand activities (predict_ligand_activities) in combination with the expression of the target genes in the previously obtained gene set, and viewed the predicted target genes corresponding to these ligands and their ranking. In addition, the above main analysis process of Nichenet can also be completed by the function nichenet_seuratobj_aggregate.

### Cytokine correlation analyses

We first obtained some cytokine ligand genes and then performed a preliminary screening and retained them if their average expression after normalization in single or multiple stages was greater than 0.25. We obtained stage-specific cytokines with significantly higher expression in one stage than in other stages and a fold change ≥ 1.05. In addition, we artificially retained some cytokines with certain expression in certain cell types, and finally, we retained 29 cytokine ligand genes. We then performed hierarchical clustering point mapping of cytokine ligand genes using Seurat, ComplexHeatmap (version 2.11.1), circlize (version 0.4.14), presto (version 1.0.0), and tidyverse (version 1.3.1) in the R (version 4.0.5) environment. We performed k-means hierarchical clustering of rows with behavioral genes, listed as stages, with the number of clusters set at 4. After obtaining stage-specific cytokine expression profiles, we used the same gene order to demonstrate their expression in different cell types using the DotPlot function. For the statistics of cell-to-cell interactions, this study used the CellTalkDB database^28^ to obtain the receptors corresponding to cytokine ligands. By defining expression greater than 0 as expressing that ligand gene or receptor gene, we obtained cell types with expression ratios greater than or equal to 50% in each stage. The potential receptor/ligand pairs defined by the study are relatively simple, i.e., there are both A (ligand cell) expressing this ligand gene A and B (receptor cell) expressing the corresponding receptor gene B in that stage, and we consider that there are potential receptor/ligand pairs gene A-gene B between A and B. Finally, the study uses igraph (version 1.3.1) for the four stages of intercellular interactions. We also screened the receptor/ligand pairs by the number of ligand-recipient pairs between a certain ligand cell and a certain receptor cell as the weight, and when the number of receptor/ligand pairs greater than or equal to the average of all weights was greater than or equal to 20, the receptor/ligand interaction was displayed. If less than 20 and the number of unscreened receptor/ligand cell pairs is greater than or equal to 20, the top 20 are selected for display in order of weight, or all are displayed if the number of unscreened receptor/ligand pairs is less than 20.

## Results

### An atlas of endometrial cells throughout the reproductive cycle in healthy women

To investigate the changes in the composition and transcriptome of endometrial immune cell populations throughout the reproductive cycle, we collected seven publicly available scRNA-seq datasets obtained from the endometrial samples of 52 total healthy women ^1,5,13,15,16,18,19^. Specifically, samples from the proliferative (n = 4) and secretory (n = 11) stages were obtained on days 1-14 and 15-28 of the menstrual cycle^1,16^, respectively. The average pregnancy duration of early pregnancy samples (n = 28) varied from 6 to 14 gestational weeks^5,13,18,19^. Likewise, the pregnancy duration of late pregnancy samples (n = 9) ranged from 33 to 40 gestational weeks (Fig. 1A)^15^.

**Fig. 1.**
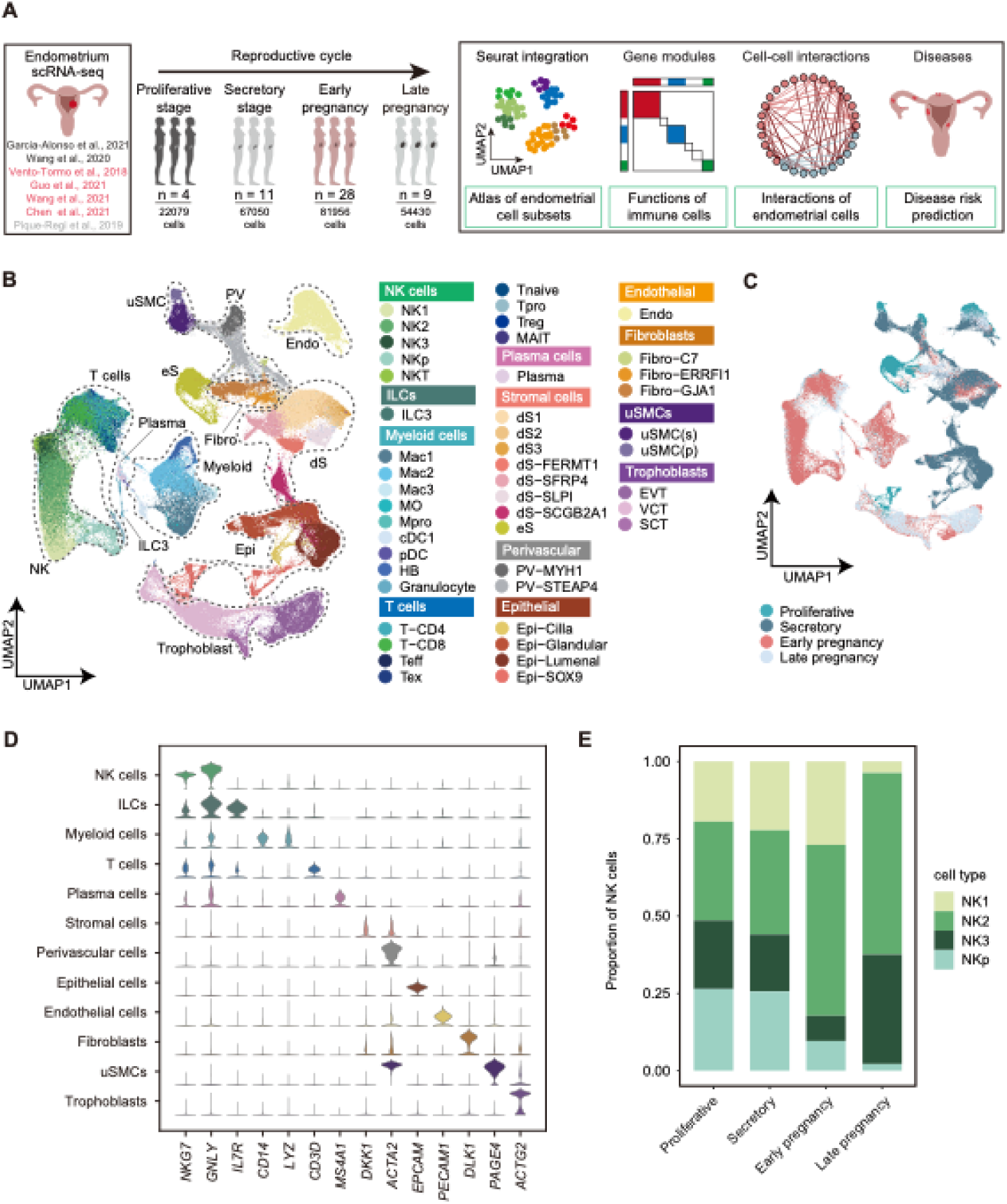
Single-cell transcriptional landscape of four stages of the reproductive cycle. **A.** A schematic outline depicting the workflow for data collection from published literature and subsequent integrated analysis. The number of samples and the number of single-cell transcriptomes collected in each stage (proliferative, secretory, early pregnancy, and late pregnancy stages) are indicated. **B.** Uniform manifold approximation and projection (UMAP) embeddings of integrated single-cell transcriptomes of four stages of samples. Cells are colored by cell subsets, and dashed circles indicate the major cell types. NK, natural killer cells; ILC, innate lymphocyte cells; p/pro, proliferative; Mac, macrophages; Mac3, maternal macrophages; HB, Hofbauer cells; DC, dendritic cells; T, T cells; Teff, effector T cells; Tex, exhausted T cells; MAIT, Mucosal-associated invariant T cells; dS, decidual stromal cells; eS, endometrial stromal cells; PV, perivascular cells; Epi, epithelial cells; Endo, endothelial cells; Fibro, fibroblasts; uSMC, uterine smooth muscle cells; SCT, syncytiotrophoblast; VCT, villous cytotrophoblast; EVT, extravillous trophoblast. **C.** UMAP embeddings of different stages illustrating no obvious batch effect in this integrated atlas. **D.** Violin plots of canonical markers (columns) for major cell types (rows). **E.** Bar plots of the cell subset composition of NK cells in four stages.

After removing cell doublets with DoubletFinder^21^ and filtering out low-quality cells, we applied Seurat^20^ to integrate the scRNA-seq datasets of endometrial samples from different stages of the human female reproductive cycle. This process resulted in a combined total set of 230,049 single-cell transcriptomes, including 22,115 cells obtained in the proliferative stage, 67,098 cells from the secretory stage, 83,111 cells from early pregnancy, and 57,725 cells from late pregnancy. The resulting gene expression matrix was normalized, and a subsequent hierarchical clustering analysis revealed the presence of 47 distinct clusters, which were visualized using uniform manifold approximation and projection (UMAP) plots. Cell lineages were identified based on predominant markers^13^, including three NK cell subsets (NK1, NK2, and NK3 cells), two macrophage subsets (Mac1 and Mac2 cells), three T-cell subsets (CD4^+^ T, CD8^+^ T, and Treg cells), dendritic cells (DCs), plasma cells, granulocytes, and nonimmune cells (including stromal cells, endothelial cells, fibroblasts, epithelial cells, perivascular cells, and trophoblasts) (Fig. 1B-D, and Supplementary Fig. 1A, B). In addition, this analysis also detected proliferative cell subsets, such as proliferative NK cells (NKp), proliferative macrophages (Mpro), and proliferative T cells (Tpro).

Comparison with cell types previously annotated in the literature^1,5,13^ showed remarkable consistency with the cell identities of populations in these datasets, thus confirming the validity of our cluster annotations (Supplementary Fig. 2A-C). In addition, stage-specific subsets of cells in our atlas were preserved through the cell-typing process. For example, endometrial stromal (eS) cells largely appeared in the proliferative stage, while decidual stromal (dS) cells only appeared in the secretory stage, early pregnancy, and late pregnancy (Fig. 1B, C and Supplementary Fig. 3A), which aligns well with the known time of differentiation from eS cells to dS cells^29^. Trophoblasts were also detected only in early and late pregnancy since they develop to initiate the invasion process after embryo implantation (Fig. 1B, C and Supplementary Fig. 3A). These collective results indicated that the single-cell transcriptome atlas of human endometrial cells from different stages of the reproductive cycle was reliable for further analysis of changes in immune cell composition, transcriptome, and intercellular interactions.

### Dynamics in the proportion of immune cells across different reproductive cycle stages

The immune cells found in the endometrium play crucial roles in facilitating a successful pregnancy, including maintaining immune tolerance, regulating trophoblast invasion, promoting fetal growth, and fighting infections^7,30-32^. Thus, we next investigated the distribution of major immune cell subsets (NK cells, macrophages, and T cells) throughout the reproductive cycle. Notably, during early pregnancy, these cell subsets all increased significantly (Supplementary Fig. 3B), indicating their importance in facilitating normal pregnancy, particularly implantation.

We further investigated the compositional changes in specific cell subsets within each major immune cell subset. We found that NK1 and NK2 cells increase during early pregnancy (Fig. 1E, and Supplementary Fig. 3C), suggesting their potential to bind to HLA class I molecules of EVTs and modulate invasion during this stage. In contrast, NK3 cells, that secrete high levels of cytokines, increase in number during late pregnancy and maintain a high proportion with NK2 cells (Fig. 1E, and Supplementary Fig. 3C). These findings were corroborated in Whettlock’s study using flow cytometry^33^. Additionally, CD4^+^ T cells were observed to increase significantly during late pregnancy (Supplementary Fig. 3D). Regarding macrophages, the proportion of Mac1 with pro-inflammatory polarization characteristics is higher during secretory and late pregnancy stages than Mac2, which exhibits anti-inflammatory polarization characteristics (Supplementary Fig. 4A-C). This is in line with overall inflammatory events that have been reported in healthy pregnancies^3^. Therefore, macrophages may play an essential role in regulating the inflammatory environment of the endometrium.

### Gene module analyses reveal the functional states and cell origin of NK cells at different stages

As the most abundant immune cells in pregnancy^3^, NK cells are the primary subject of our investigation into functional changes throughout the reproductive cycle. To determine the gene expression differences between NK cell subsets, we utilized consensus nonnegative matrix factorization (cNMF) and hierarchical clustering strategies, resulting in the identification of 8 distinct stage-specific gene modules (GMs) (see Methods). Expression changes in each of these GMs during different stages of pregnancy were found to be similar amongst the various NK cell subsets (Fig. 2A, B, and Supplementary Fig. 5). Within GM1, genes involved in ATP metabolism (glycolysis: *TPI1*, *GAPDH*; ATP synthesis and catabolism: *ATP5PF*, *ATP5MF*, *ATP5MC2*, *ATP5MG*; aerobic respiration: *UQCRQ*, *COX7A2*) were found to be highly expressed in NK cells during early pregnancy (Fig. 2B, C, and Supplementary Fig. 6, 7A). Our findings thus supported previous work which characterized the active glycolytic metabolism amongst NK1 cells within early pregnancy^13^ and furthermore, revealed that oxidative phosphorylation metabolism (resulting in the production of more ATP than glycolytic and TCA cycle metabolism) of NK cells is more active during early pregnancy than in other stages, as determined using scMetabolism^34^ (Supplementary Fig. 7B). Taken together, our results suggest an active functional state of NK cells during early pregnancy.

**Fig. 2.**
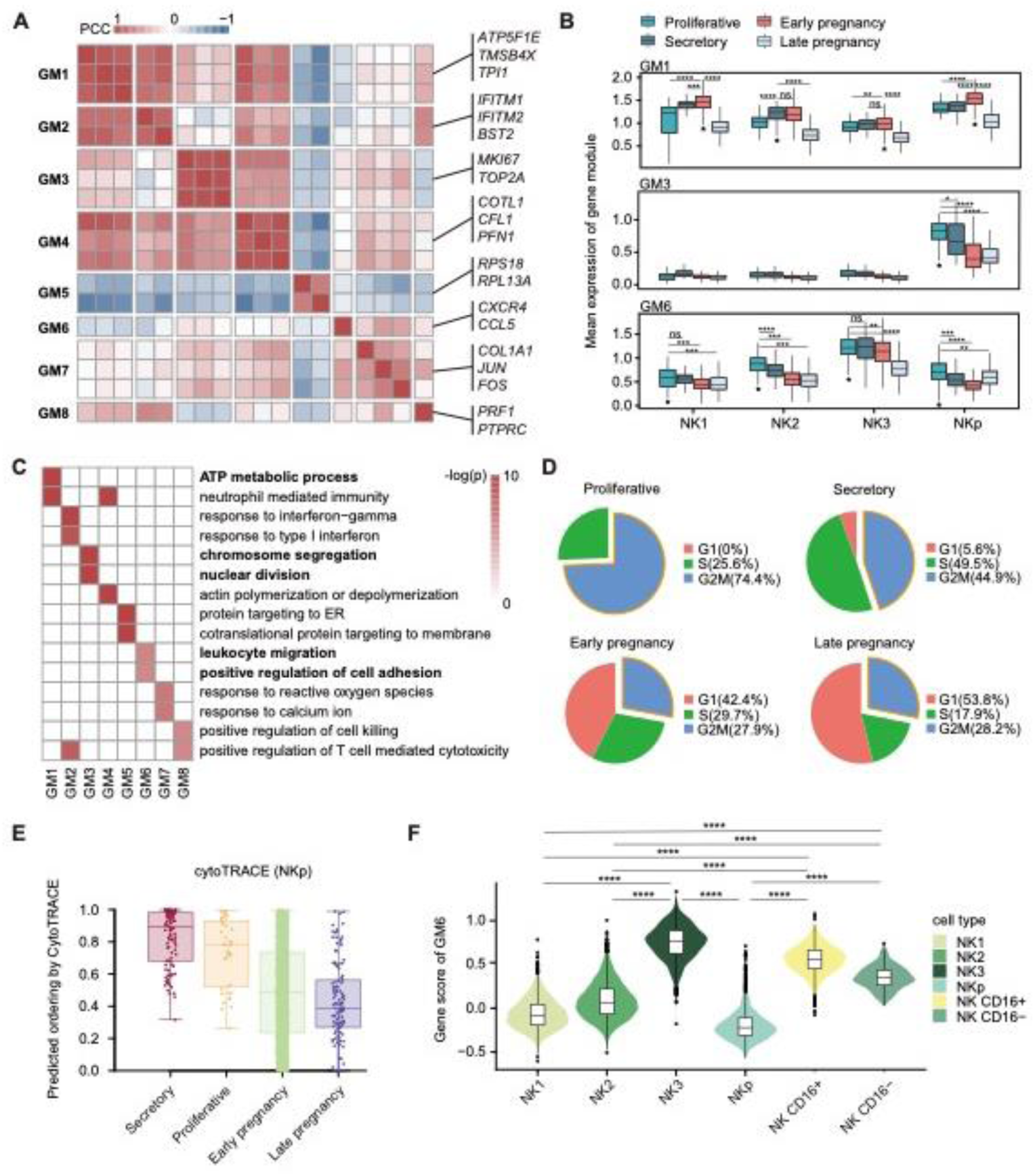
Transcriptome changes in NK cells in four stages. **A.** Heatmap of Pearson correlation coefficients (PCCs) of the relative normalized gene expression of modules from Consensus nonnegative matrix factorization (cNMF). **B.** Box plots of the relative normalized gene expression of GM1, GM3, and GM6 in different NK cell subsets in four stages. **C.** Gene Ontology (GO) analysis of stage-specific gene modules. **D.** Pie plots show the proportion of NKp cells in different cell cycle phases in four stages. **E.** Box plots of NKp differentiation potential calculated by cytoTRACE in four stages. **F.** Violin plots of GM6 gene scores of endometrial and peripheral blood NK cell subsets. Statistical significance between different stages was evaluated with the Wilcoxon rank-sum test (two-tailed). In B) and F), the mean and interquartile range (IQR), with whiskers extending to 1.5×IQR, are shown in the plots. **** P < 0.0001, *** P < 0.001, ** P < 0.01, * P <0.05.

Beyond this, the origin of NK cells remains unclear ^7^, leading us to focus on GM3 (mitosis: *MKI67*, *TOP2A, etc.*) and GM6 (leukocyte migration: *CCL5*, *CXCR4*, *etc.*), both of which were observed to be highly expressed during the menstrual cycle (Fig. 2B, C). We found that GM3 was highly expressed in NKp, suggesting that NKp has the strongest proliferative capacity in the menstrual cycle (Fig. 2B), which was further confirmed by calculating cell cycle phase scores based on canonical markers^35^ (Fig. 2D). Our previous work demonstrated that NKp has the potential to differentiate into other NK cell subsets in early pregnancy^5^. To further elucidate this phenomenon, we applied cytoTRACE to NK cells of four stages, and discovered that NKp from different stages showed similar differentiation programs (Supplementary Fig. 8A-D). Furthermore, we found that NKp of non-pregnant stages had a higher differentiation potential compared to NKp of pregnancy (Fig. 2E). Meanwhile, the proportion of NKp decreased gradually as the pregnancy progresses (Fig. 1E). These results suggest that NKp of non-pregnant stages may have a greater proliferative and differentiation capacity than NKp of pregnancy.

GM6 was enriched for functions related to leukocyte migration, and given the possibility that peripheral blood NK (pbNK) cells may act as a source of endometrial NK cells^36^, we assessed the expression levels of GM6 genes in endometrial NK cells. We discovered that the NK3 subset exhibited strong gene expression (*e.g.*, *CXCR4*) related to chemotactic properties, akin to pbNK cells, and especially CD16^+^ pbNK cells (Fig. 2F, and Supplementary Fig. 9A). Moreover, *CXCL12* (the ligand of *CXCR4*) was found to have high expression levels in eS cells and fibroblasts during non-pregnant stages (Supplementary Fig. 9B), thereby suggesting that NK3 cells may be recruited by these cells.

### Endometrial immune cells display a strong IFN-γ response in early pregnancy

We also conducted GM analysis on both macrophages and T cells. We discovered that similar to NK cells, both macrophages and T cells exhibit a potent type II interferon (IFN-γ) response in early pregnancy (macrophage: GM5, T cell: GM1) (Supplementary Fig. 10, and Supplementary Fig. 11). By using CytoSig^26^ to predict cytokine signaling activity, we further substantiated this observation (Fig. 3A). Further investigation on the genes contained in these interferon-associated GMs, we discovered that during early pregnancy, Mac1 and Mac2 express chemokines such as *CCL3* and *CCL4* (Fig. 3B). Additionally, NK cells and T cells display elevated levels of MHC class I molecules, while macrophages display increased levels of MHC class II molecules (Fig. 3B). These findings suggest that immune cells exhibited an activated state and potential antiviral functions^37^ during early pregnancy.

**Fig. 3.**
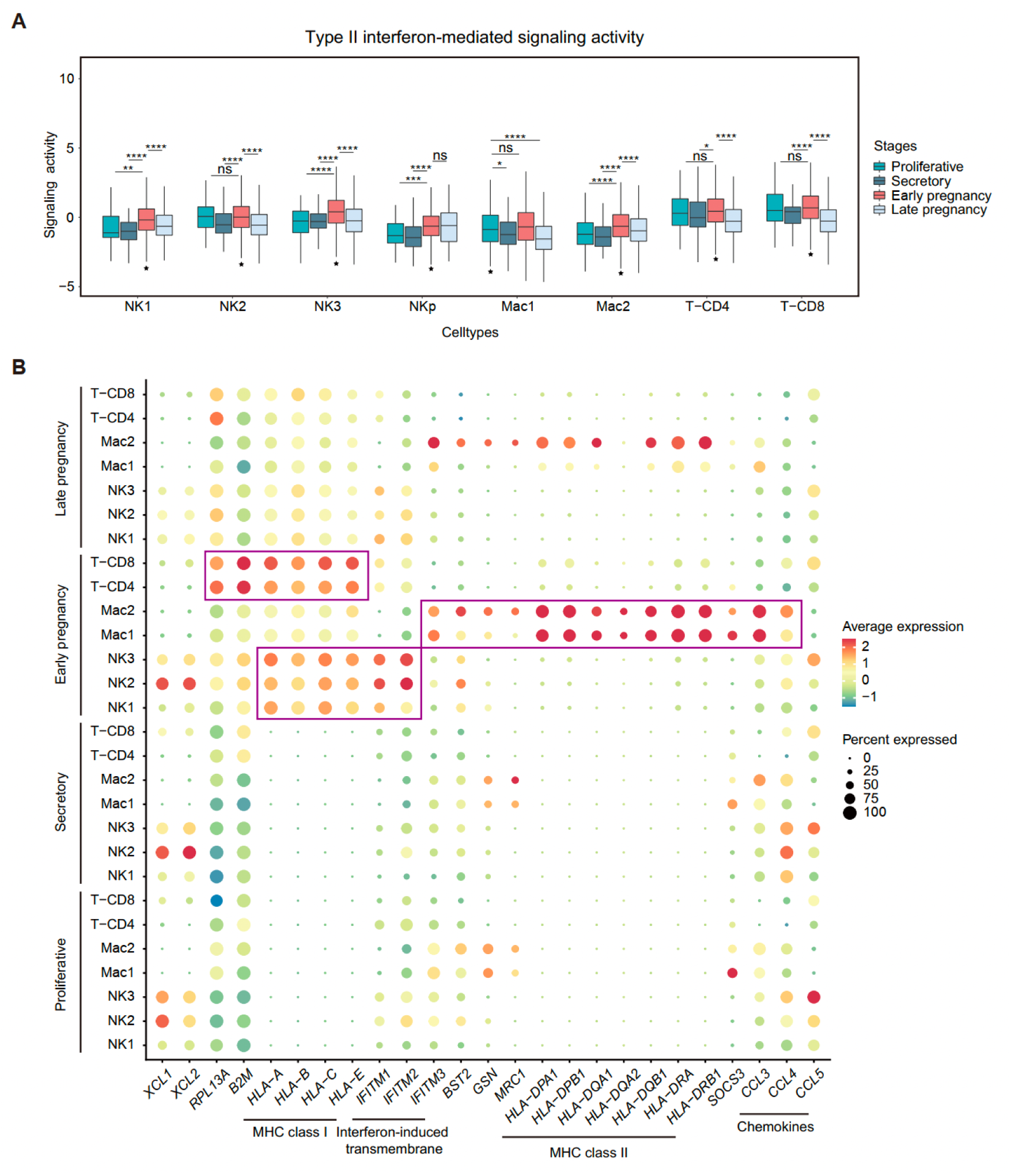
Type II interferon response of major immune cell subsets. **A.** Box plots predicting the type II interferon signaling activity of major immune cell subsets in four stages. **B.** Dot plots of the expression of type II interferon response genes of major immune cell subsets in four stages. Statistical significance between different stages was evaluated with the Wilcoxon rank-sum test (two-tailed). In A), the mean and interquartile range (IQR), with whiskers extending to 1.5×IQR, are shown in box plots. **** P < 0.0001, *** P < 0.001, ** P < 0.01, * P <0.05.

### Stage-specific cell-cell interactions in the menstrual cycle promote cell proliferation and decidualization

Given that cytokines and chemokines can mediate cell-cell interactions at the maternal-fetal interface that play a fundamental role in pregnancy success^13,38,39^, we next examined cell-cell interactions involving these signaling molecules at different stages of the reproductive cycle. Initially, we generated a set of stage-specific cytokines, with approximately 93% of them encoding secreted proteins (Supplementary Fig. 11A). We utilized CellTalkDB^28^ to obtain their respective receptors. Potential stage-specific receptor/ligand pairs were then predicted by analyzing their expression in different cell subsets during different stages (see Methods). Among the four stages, the late pregnancy exhibited the highest diversity of ligand-receptor pairs (n = 20 receptor/ligand pairs) as predicted, followed by early pregnancy (n = 19) and the proliferative stage (n = 12), while the secretory stage had the fewest predicted pairs (n = 5) (Fig. 4A, and Supplementary Fig. 11B).

**Fig. 4.**
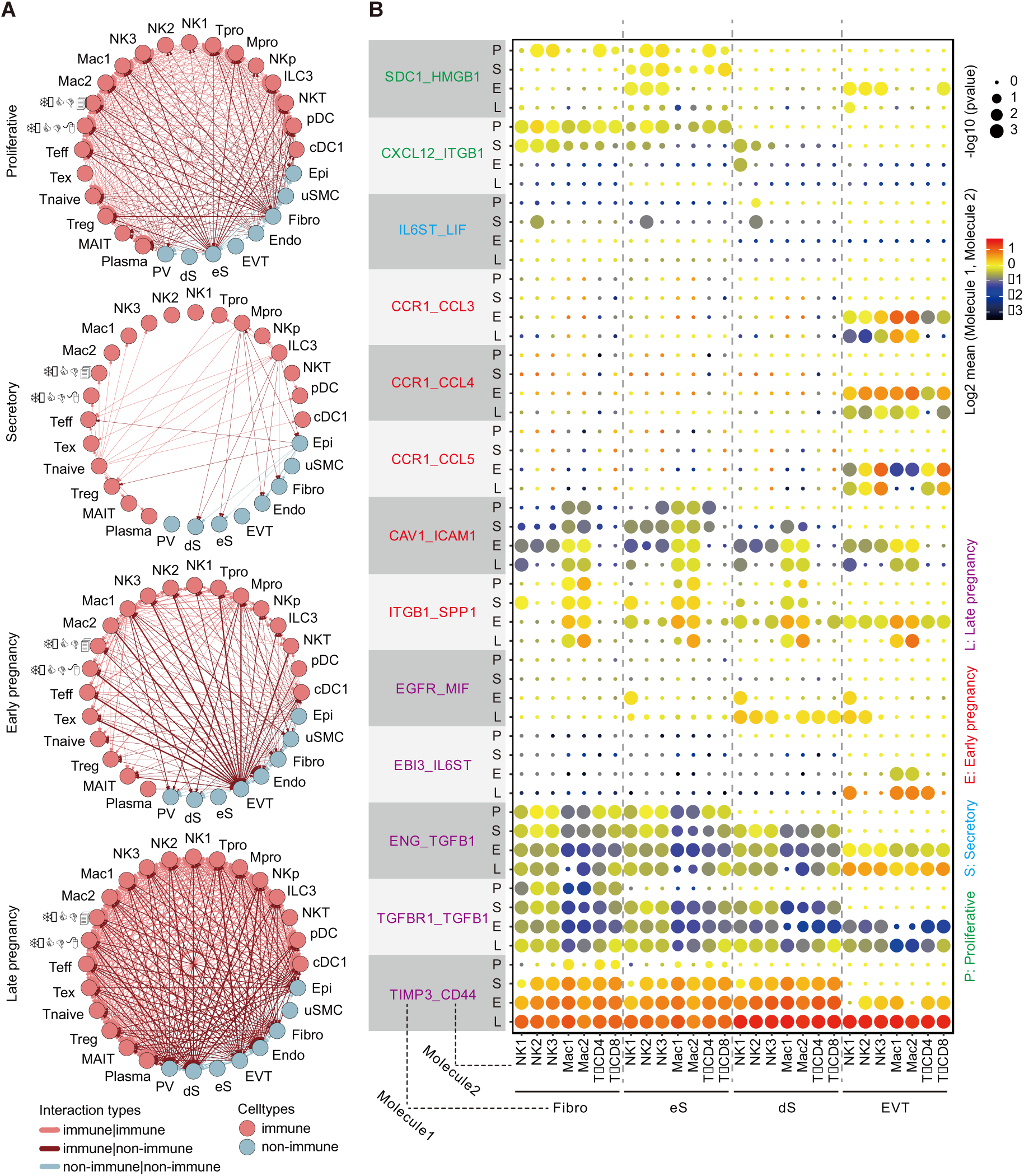
Cell-cell interactions of the endometrium in four stages. **A.** Circos plots of the receptor/ligand pairs between different cell subsets in four stages. **B.** Dot plots of the expression of receptor/ligand pairs between different cell subset pairs.

We utilized CellPhoneDB^40^ to explore the specific cell-cell interactions between non-immune and immune cell during each stage (Fig. 4B). During the proliferative stage, nearly all endometrial cell types, including immune cells, exhibited elevated levels of *HMGB1*, promoting cell proliferation. *SDC1* (receptor of *HMGB1*) was highly expressed by eS cells and fibroblasts, which may lead to the endometrial thickening in the proliferative stage (Fig. 4B). In addition, eS cells and fibroblasts express ligands involved in the CXCL signaling pathway in the proliferative stage, and the major immune subsets express their corresponding receptors, thus suggesting a possible mechanism for immune cell recruitment. Notably, this interaction analysis identified the expression of *LIF* in NK2 cells during the secretory stage (Fig. 4B). A previous study has shown that *LIF* can enhance eS decidualization^41^, which, in conjunction with our results, suggesting that the NK2 cells may participate in promoting decidualization.

### Stage-specific cell-cell interactions in pregnancy stages promote tissue remodeling and immune homeostasis at the maternal-fetal interface

Upon investigating the potential ligand-receptor pairs during early pregnancy, we found that major immune cell subsets highly expressed *CCL3*, *CCL4* and *CCL5*, while EVTs specifically expressed their receptors (*CCR1*, *SDC4*, and *ACKR2*). This may facilitate EVT invasion in early pregnancy and EVT may also regulate chemokine levels to prevent excessive invasion of itself. In addition, non-immune cells exhibited similar patterns in the expression of receptors of *ICAM1* and *SPP1*. The former participates in leukocyte-endothelial cell adhesion^42^, while the latter binds integrins to promote cell adhesion, migration, and differentiation via cell-cell and cell-extracellular matrix interactions^43^. Our results showed that ligands of both receptors were highly expressed in Mac1 and Mac2 in early pregnancy (Fig. 4B), suggesting their potential involvement in the reconstruction of the endometrial immune microenvironment.

During late pregnancy, our findings showed that the receptor of *MIF* was expressed at a high level in dS cells and EVTs (Fig. 4B). MIF is a pro-inflammatory cytokine^3,44^ that can bind to the human epidermal growth factor receptor (EGFR), thus preventing its activation. According to our results, this can lead to immune cells using *MIF* to hinder the proliferation of dS cells in late pregnancy. *EBI3*, an immunomodulator involved in regulating NK cells^45^, was highly expressed in EVTs, while its receptor *IL6ST* was highly expressed in NK1, Mac1, Mac2, and CD4^+^ T cell subsets (Fig. 4B, and Supplementary Fig. 11A). Moreover, our analysis showed that *TGFB1*, which can reduce trophoblast invasiveness^46^, was highly expressed in NK cells and T cells (Fig. 4B). The receptors of *TGFB1* (*ENG* and *TGFBR1*) were expressed by EVT. EVT invasion may be also influenced by tissue inhibitor of metalloproteinase 3 (*TIMP3*), which was highly expressed in several non-immune cell subsets (*i.e.*, perivascular cells, stromal cells, endothelial cells, fibroblasts, epithelial cells, and trophoblasts) (Fig. 4B, and Supplementary Fig. 11A), and may limit excessive EVT invasion through the suppression of the metalloproteinase-mediated extracellular matrix remodeling process.

### Association analysis with disease risk genes reveals the potential clinical significance of stage-specific genes in preeclampsia

Next, we evaluated the clinical relevance of stage-specific gene expression profiles in immune cells. Specifically, we obtained stage-specific genes from previous GM analyses and collected risk gene sets for five common reproductive diseases from the Harmonizome database^47^, including endometriosis, implantation failure, gestational diabetes (GD), RPL, and preeclampsia (Supplementary Table 1). We then established enrichment scores through Fisher’s exact test, which measured the association between the risk genes and the stage-specific genes.

We first found that RPL showed the highest enrichment score during early pregnancy in immune cells (Fig. 5A). Upon analyzing the data, we discovered that risk genes pertaining to interactions (*e.g.*, *HLA-A*, *HLA-DRB1*, and *TNF*), all of which were previously identified in other studies^5,18^, were noticeably enriched in early pregnancy associated with RPL (Fig. 5A). Interestingly, our results showed that these genes may also be associated with other reproductive diseases except for implantation failure (Fig. 5A). Additionally, we linked the upregulated genes of major immune cell subsets specific to early pregnancy obtained from scRNA-seq disease data collected from our previous research^5^ to the enriched stage-specific risk genes affiliated with RPL. Our results demonstrated a significant association between the enriched stage-specific risk genes in RPL patients and the upregulated genes pertaining to major immune cell subsets of early pregnancy (Supplementary Fig. 12A-C). Furthermore, we noticed that several specific interaction-related genes linked to various stages of pregnancy (*TGFB1* for late pregnancy, *ICAM1* and *CCL5* for early pregnancy) were differentially enriched in endometriosis and preeclampsia. The results suggested that immune-associated stage-specific interactions are vital for healthy pregnancy while also suggesting a high risk of being associated with reproductive diseases.

**Fig. 5.**
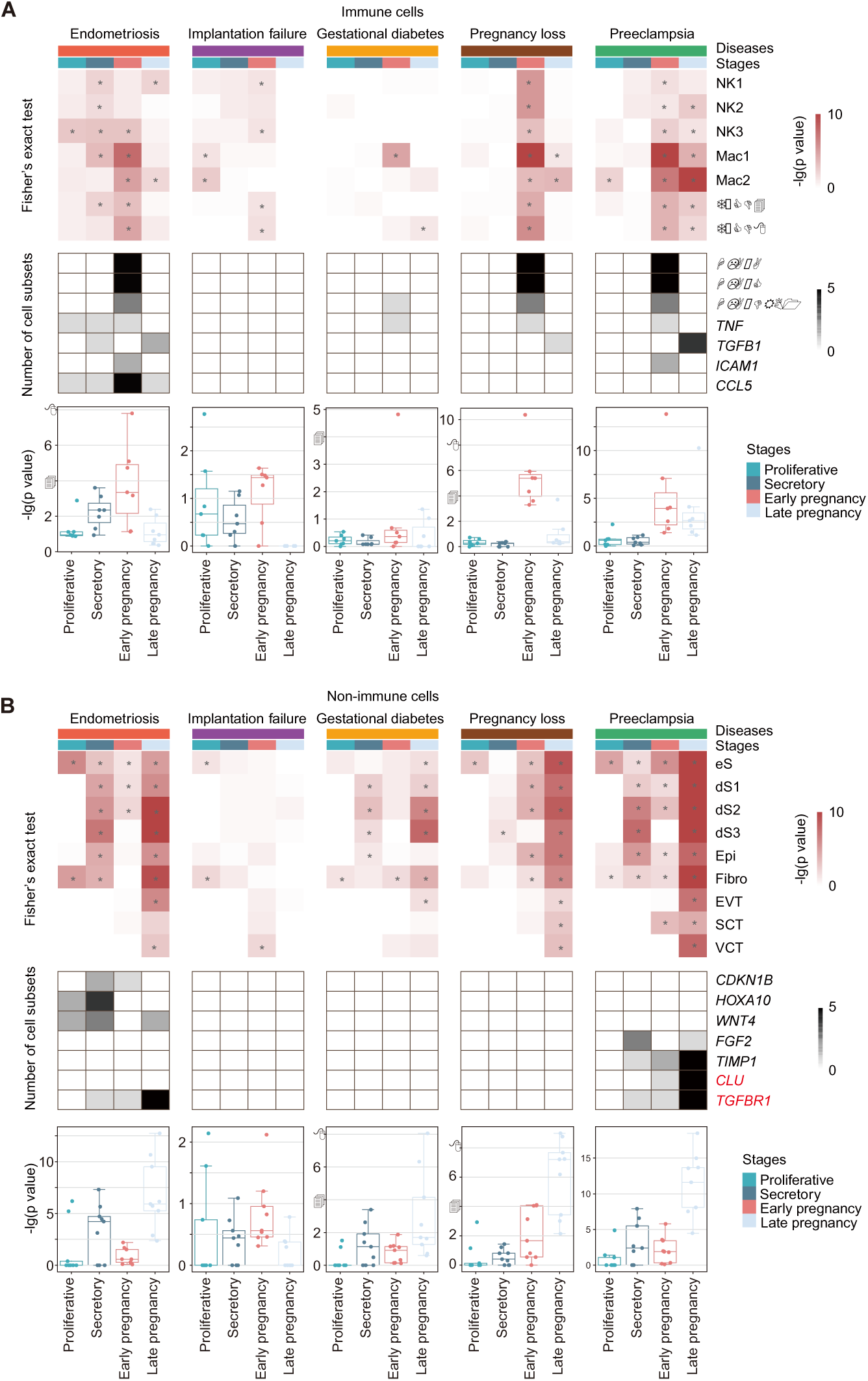
Clinical relevance between the reproductive cycle and reproductive diseases. **A, B.** Heatmaps of the enrichment score of disease risk genes within stage-specific genes of immune (A) or nonimmune (B) cell subsets (top), heatmaps of the number of occurrences of risk genes of endometrial cell subsets in different reproductive cycle stages (middle), and box plots of the enrichment score of immune/nonimmune cell subsets in different reproductive cycle stages (bottom). -log (P value) was defined as the enrichment score. * P <0.05.

We also evaluated the clinical relevance of stage-specific gene expression profiles in non-immune cells (Fig. 5B). Notably, certain genes that are considered pivotal for the appearance of endometriosis, for instance, *CDKN1B*^48^, *HOXA10*^49^, and *WNT4*^50^, were found to be enriched in the secretory stage (Fig. 5B). *HOXA10* and *WNT4* were also detected to be enriched in the proliferative stage (Fig. 5B), implying that they might be associated with different stages of endometriosis development. By performing association analysis with preeclampsia risk genes, we revealed the clinical relevance of stage-specific genes like *FGF2*, *TIMP1*, *CLU*, and *TGFBR1* from secretory, early pregnancy, and late pregnancy stages (Fig. 5B). These genes were reported to be associated with the treatment or severity of preeclampsia^51-54^. Our results showed several genes with high enrichment scores in at least one cell subset, such as, *FGF2* (secretory stage and late pregnancy), *TIMP1* (secretory stage, early and late pregnancy)*, CLU* (early and late pregnancy) and *TGFBR1* (secretory stage, early and late pregnancy) (Fig. 5B). Although symptoms of preeclampsia typically occur after the 20^th^ week of gestation^47^, our results suggested the possibility of early detection of preeclampsia at the transcriptome level.

In a recent study, maternal blood samples were utilized to identify and verify cfRNA transcriptomic changes that are linked to preeclampsia, and genes that can differentiate between patients with preeclampsia and healthy individuals in early pregnancy were identified^55^ (Supplementary Table 2). We performed Fisher’s exact test on our stage-specific genes and these cfRNA markers to explore the association between them. We found that the enrichment score of non-immune cells was high in all four stages and that of immune cells was high in the secretory stage (Supplementary Fig. 13A, B), suggesting that the endometrium and peripheral blood show a synergistic relationship in preeclampsia development. By computing intersections of risk genes, cfRNA genes, and stage-specific genes, two candidate risk genes (*CLU* and *TGFBR1*) were identified, especially *TGFBR1,* which was highly expressed in both immune cells and non-immune cells during secretory phase (Supplementary Fig. 13C, D). This demonstrated that cfRNA can be utilized as an early indicator for peripheral screening for preeclampsia. In general, our integrated analysis suggests that there could be potential benefits to utilizing the genes’ signatures unique to each stage for early screening of reproductive diseases.

## Discussion

Immune cell infiltration of the endometrium has been proposed as an essential step for promoting a successful pregnancy and immune homeostasis^4^. The large majority of published studies have used scRNA-seq to examine endometrial cell heterogeneity exclusively in pregnancy or nonpregnancy stages. In this study, we compiled scRNA-seq datasets from several cohorts obtained at four distinct stages of the reproductive cycle, including the proliferative, secretory, early pregnancy, and late pregnancy stages. Through advanced bioinformatic analyses, we characterized the immune landscape at each stage to identify changes in the composition, function, and cell-cell interactions of endometrial cell populations (Fig. 6). Analyses of proportions revealed changes in the proportions of NK1 and NK2 subsets during four stages of reproductive cycle differ from those observed in NK3 cells. Identification of stage-specific GMs revealed that these immune cells have a strong type II interferon response in early pregnancy. Additionally, we found cell-cell interactions related to proliferation promotion in the proliferative stage and tissue remodeling in early pregnancy. We also identified *CLU* and *TGFBR1* as potential biomarkers for the early detection of preeclampsia. Our integrative analysis of immune cell subsets during the reproductive cycle provided insights into the advancement of reproductive processes.

**Fig. 6.**
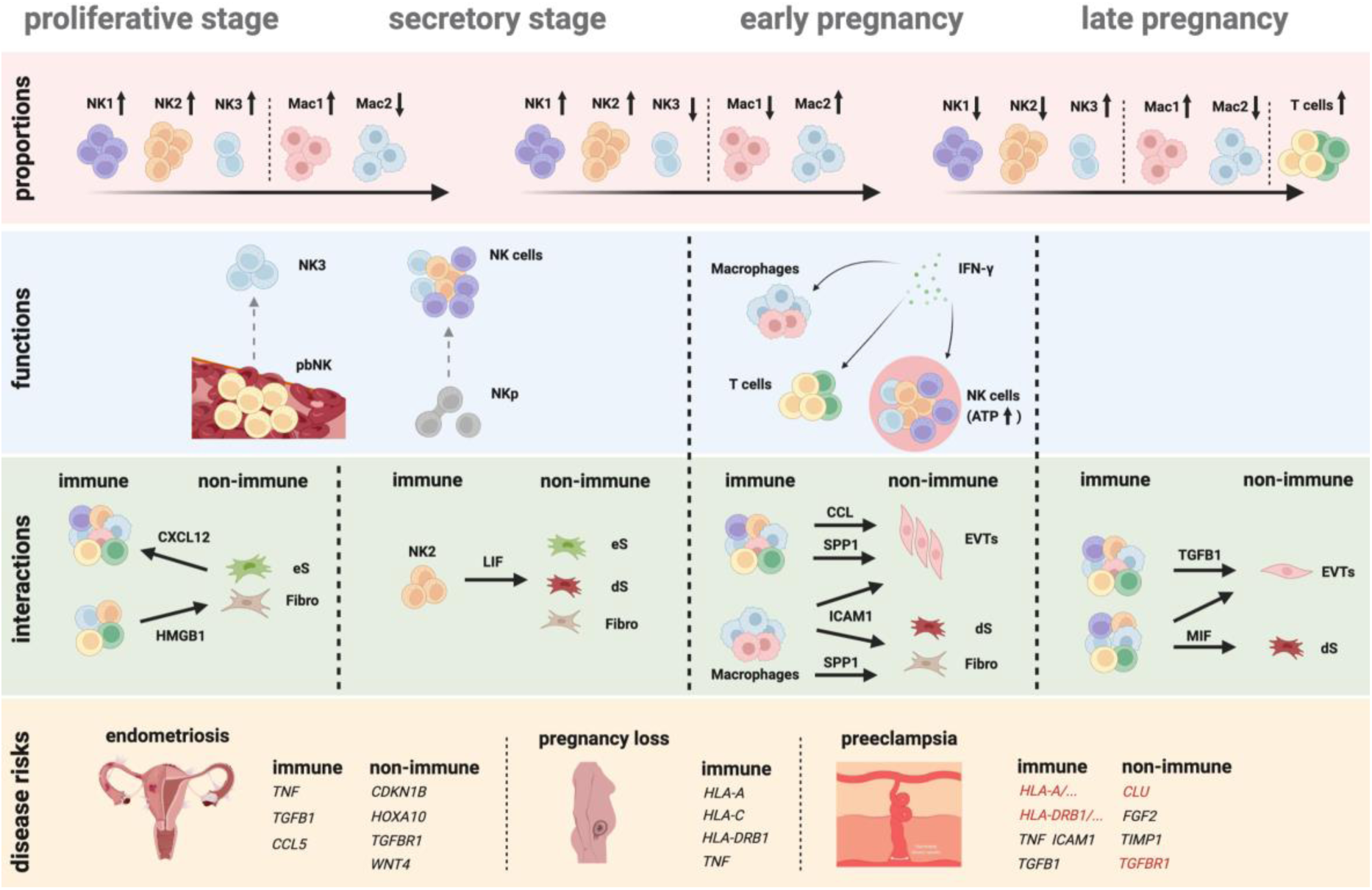
Summary graph. Advanced bioinformatic analyses was implemented to investigate integrated scRNA data. For cell proportions, we primarily observed alterations in the proportions of NK1, NK2, NK3, Mac1, Mac2, and T cells across four stages. For functions, we mainly found that NKp cells have a strong proliferative capacity and differentiation potential during non-pregnant stages. NK3 displayed strong gene expression that is associated with chemotactic properties, similar to pbNK cells. NK cells display high oxidative phosphorylation metabolism activity during early pregnancy, and in conjunction with macrophages and T cells, exhibit a potent type II interferon response. For cell-cell interactions, we identified ligand-receptor pairs that are specific to each stage of reproductive cycle, for instance, the proliferative stage was associated with promotion of cell proliferation, the secretory stage with promotion of stromal cell decidualization, early pregnancy with promotion of EVTs invasion, and late pregnancy with inhibition of proliferation and EVTs invasion. For disease risks, we identified numerous stage-specific risk genes and in combination with cfRNA identified potential predictors associated with preeclampsia such as *CLU* and *TGFBR1*, as well as genes that may play a pivotal role in a variety of reproductive diseases in our results such as HLA class I/II molecules.

Decidual NK cells have been thought to be derived from CD34^+^ hematopoietic stem cells, mature from immature endometrial NK cells^38,56,57^, or to migrate from peripheral blood NK cells^7,36^. We found that NKp cells have a strong proliferative capacity and differentiation potential during non-pregnant stages, and this signal diminishes during pregnancy. The uterus maintains a low oxygen concentration in the early stages of implantation, while appearing to rise in early and mid-pregnancy^58^. We speculate that low oxygen concentrations during non-pregnant stages favor glycolytic metabolism and proliferation of NKp^59^, whereas oxygen concentrations during early pregnancy favor a strong oxidative phosphorylation metabolism and functions of NK cells. We also observed that the relative proportions of NK1 and NK2 cells showed an increase in early pregnancy and a decrease in late pregnancy. Both existing studies^5,13,18^ and our results show that NK1 and NK2 cells have similar transcriptome profiles compared with NKp cells. In contrast, NK3 cells have the least transcriptome similarity to NKp cells, while they exhibit a chemokine pattern similar to that of pbNK cells. Several studies of decidua in mid- and late pregnancy failed to separate uterine leukocytes from maternal or fetal blood leukocytes^60^, indicating that uterine leukocytes are similar to peripheral blood leukocytes at this time, which may be associated with an increase in the proportion of NK3 in late pregnancy. Thus, we speculate that NK cells may derive from NKp and pbNK cells, while NK1/2 and NK3 may arise from different precursor cells.

During the process of placentation, EVT migrates into the maternal uterus to remodel spiral arteries^13^. We identified the cytokines *CCL3*, *CCL4*, *CCL5*, *ICAM1*, and *SPP1* during early pregnancy, which may help to promote EVT invasion. SPP1 was previously reported to be produced by NK cells and macrophages^13,60^, and its expression was higher in macrophages than in NK cells. Excessive invasion of EVT is harmful, and the rate of EVT invasion decreases during pregnancy^61^. TGFB1 and TIMP3 were identified during late pregnancy in our results, which are both reported to regulate EVT invasion^61^. Immune cells (NK cells, macrophages, T cells, and DCs) and non-immune cells (fibroblasts, endothelial cells, stromal cells, and other trophoblast cells) appear to be involved in this process. Mast cells and neutrophils may also regulate EVT invasion^61^.

Preeclampsia stands as one of the most profound complications of pregnancy and the early diagnosis of this affliction can augment its prognosis. Although researchers have utilized proteomics and transcriptomics to identify potential biomarkers^62,63^, a dependable screening test for its development remains elusive^64^. Notably, a recent study showed that preeclampsia could be distinguished and predicted well during early pregnancy (≤12 weeks gestation) by using peripheral blood cfRNA genes^55^. These nucleic acids in the maternal blood may be released from the uterus^64^. In this study, we also identified two risk genes, *CLU* and *TGFBR1*, which were reported in the aforementioned cfRNA study. According to the two-stage theory, preeclampsia is associated with placental insufficiency and endothelial dysfunction^64^. Indeed, stage-specific risk genes in our results (*e.g.*, *CLU*, *TGFBR1*, *FGF2*, and *TIMP1*) were found to be associated with the protection of the normal function of vascular endothelial cells^65-69^. Since FGF2, TIMP1 and TGFBR1 appeared in the secretory stage, before the appearance of the placenta, we speculate on the prediction of preeclampsia from the endometrium and even peripheral blood during the secretory stage.

There are some limitations in our analyses. Firstly, our analysis shows only the changes that characterize the different stages of the reproductive cycle and not the real continuous changes. During the main reproductive cycle stages, we also lack data on healthy pregnant women during mid-pregnancy. Secondly, the proportion of immune cells is relatively low in the menstrual cycle, and although this has been described^7^, it may have some impact on what we have found in the menstrual cycle. Nevertheless, our new findings in this study provide a reference for further investigation of the multifaceted changes in immune cells in different stages of the healthy female reproductive cycle in the future and provide clues for the prediction and treatment of pregnancy-related diseases.

## Data Availability Statement

Our expression data for different stages are also available online from published studies^1,5,13,15,16,18,19^. They are available at ArrayExpress (E-MTAB-6701, and E-MTAB-10287), dbGaP (phs001886.v1.p1), GSA (CRA002181, and HRA000237) and GEO (GSE164449, and GSE111976).

## Acknowledgements

**Funding**: This work was supported by the National Natural Science Foundation of China grants (32270978 to C.G.), We thank the USTC supercomputing center and the School of Life Science Bioinformatics Center for providing computing resources for this project.

## Author Contributions

C.G. conceived and supervised the project. K.C. and Q.Y. collected public data and performed data analysis. C.G., K.C., and Q.Y. interpreted data with help from Q.S., J.W., B.F., X.S., and X.L.. J.F. contributed to the revision of the manuscript. K.C., C.G., and Q.Y. wrote the manuscript with help from all the other authors. All authors reviewed the manuscript and consented for publication.

## References

1. Garcia-Alonso L, Handfield LF, Roberts K, et al. Mapping the temporal and spatial dynamics of the human endometrium in vivo and in vitro. Nat Genet 2021; 53(12): 1698–711.

2. Soma-Pillay P, Nelson-Piercy C, Tolppanen H, Mebazaa A. Physiological changes in pregnancy. Cardiovasc J Afr 2016; 27(2): 89–94.

3. Zhang X, Wei H. Role of Decidual Natural Killer Cells in Human Pregnancy and Related Pregnancy Complications. Front Immunol 2021; 12: 728291-.

4. Arck PC, Hecher K. Fetomaternal immune cross-talk and its consequences for maternal and offspring’s health. Nat Med 2013; 19(5): 548–56.

5. Guo C, Cai P, Jin L, et al. Single-cell profiling of the human decidual immune microenvironment in patients with recurrent pregnancy loss. Cell Discov 2021; 7(1): 1–16.

6. Bulmer JN, Morrison L, Longfellow M, Ritson A, and Pace D. Granulated lymphocytes in human endometrium: histochemical and immunohistochemical studies. Human Reproduction 1991; 6(6): 791–8.

7. Yang F, Zheng Q, Jin L. Dynamic Function and Composition Changes of Immune Cells During Normal and Pathological Pregnancy at the Maternal-Fetal Interface. Front Immunol 2019; 10: 2317–32.

8. Vallve-Juanico J, Houshdaran S, Giudice LC. The endometrial immune environment of women with endometriosis. Hum Reprod Update 2019; 25(5): 564–91.

9. King A. Uterine leukocytes and decidualization. Hum Reprod Update 2000; 6(1): 28–36.

10. Mselle TF, Meadows SK, Eriksson M, et al. Unique characteristics of NK cells throughout the human female reproductive tract. Clin Immunol 2007; 124(1): 69–76.

11. Tsuda S, Nakashima A, Shima T, Saito S. New Paradigm in the Role of Regulatory T Cells During Pregnancy. Front Immunol 2019; 10: 573.

12. Barrozo ER, Aagaard KM. Human placental biology at single-cell resolution: a contemporaneous review. BJOG 2022; 129(2): 208–20.

13. Vento-Tormo R, Efremova M, Botting RA, et al. Single-cell reconstruction of the early maternal-fetal interface in humans. Nature 2018; 563(7731): 347-53.

14. Liu Y, Fan X, Wang R, et al. Single-cell RNA-seq reveals the diversity of trophoblast subtypes and patterns of differentiation in the human placenta. Cell Res 2018; 28(8): 819–32.

15. Pique-Regi R, Romero R, Tarca AL, et al. Single cell transcriptional signatures of the human placenta in term and preterm parturition. Elife 2019; 8: 1–22.

16. Wang W, Vilella F, Alama P, et al. Single-cell transcriptomic atlas of the human endometrium during the menstrual cycle. Nat Med 2020; 26(10): 1644–53.

17. Huhn O, Ivarsson MA, Gardner L, et al. Distinctive phenotypes and functions of innate lymphoid cells in human decidua during early pregnancy. Nat Commun 2020; 11(1): 381.

18. Wang F, Jia W, Fan M, et al. Single-cell Immune Landscape of Human Recurrent Miscarriage. Genomics Proteomics Bioinformatics 2021; 19(2): 208–22.

19. Chen P, Zhou L, Chen J, et al. The Immune Atlas of Human Deciduas With Unexplained Recurrent Pregnancy Loss. Front Immunol 2021; 12: 689019–35.

20. Hao Y, Hao S, Andersen-Nissen E, et al. Integrated analysis of multimodal single-cell data. Cell 2021; 184(13): 3573–87.

21. McGinnis CS, Murrow LM, Gartner ZJ. DoubletFinder: Doublet Detection in Single-Cell RNA Sequencing Data Using Artificial Nearest Neighbors. Cell Syst 2019; 8(4): 329–37.

22. Korsunsky I, Millard N, Fan J, et al. Fast, sensitive and accurate integration of single-cell data with Harmony. Nat Methods 2019; 16(12): 1289–96.

23. Lopez R, Regier J, Cole MB, Jordan MI, Yosef N. Deep generative modeling for single-cell transcriptomics. Nat Methods 2018; 15(12): 1053–8.

24. Kotliar D, Veres A, Nagy MA, et al. Identifying gene expression programs of cell-type identity and cellular activity with single-cell RNA-Seq. Elife 2019; 8.

25. Yu G, Wang LG, Han Y, He QY. clusterProfiler: an R package for comparing biological themes among gene clusters. OMICS 2012; 16(5): 284–7.

26. Jiang P, Zhang Y, Ru B, et al. Systematic investigation of cytokine signaling activity at the tissue and single-cell levels. Nat Methods 2021; 18(10): 1181–91.

27. Browaeys R, Saelens W, Saeys Y. NicheNet: modeling intercellular communication by linking ligands to target genes. Nat Methods 2020; 17(2): 159–62.

28. Shao X, Liao J, Li C, Lu X, Cheng J, Fan X. CellTalkDB: a manually curated database of ligand–receptor interactions in humans and mice. Briefings in Bioinformatics 2020; 22(4).

29. Lee JY, Lee M, Lee SK. Role of endometrial immune cells in implantation. Clin Exp Reprod Med 2011; 38(3): 119–25.

30. Liu S, Diao L, Huang C, Li Y, Zeng Y, Kwak-Kim JYH. The role of decidual immune cells on human pregnancy. J Reprod Immunol 2017; 124: 44–53.

31. Hanna J, Goldman-Wohl D, Hamani Y, et al. Decidual NK cells regulate key developmental processes at the human fetal-maternal interface. Nat Med 2006; 12(9): 1065–74.

32. Fu B, Zhou Y, Ni X, et al. Natural Killer Cells Promote Fetal Development through the Secretion of Growth-Promoting Factors. Immunity 2017; 47(6): 1100–13 e6.

33. Whettlock EM, Woon EV, Cuff AO, Browne B, Johnson MR, Male V. Dynamic Changes in Uterine NK Cell Subset Frequency and Function Over the Menstrual Cycle and Pregnancy. Frontiers in Immunology 2022; 13.

34. Wu Y, Yang S, Ma J, et al. Spatiotemporal Immune Landscape of Colorectal Cancer Liver Metastasis at Single-Cell Level. Cancer Discov 2022; 12(1): 134–53.

35. Nestorowa S, Hamey FK, Pijuan Sala B, et al. A single-cell resolution map of mouse hematopoietic stem and progenitor cell differentiation. Blood 2016; 128(8): e20–31.

36. Benedikt S, Jonna B, Hanna J, et al. Continuous human uterine NK cell differentiation in response to endometrial regeneration and pregnancy. Sci Immunol 2021; 6(56): eabb7800.

37. Ranasinghe S, Lamothe PA, Soghoian DZ, et al. Antiviral CD8(+) T Cells Restricted by Human Leukocyte Antigen Class II Exist during Natural HIV Infection and Exhibit Clonal Expansion. Immunity 2016; 45(4): 917–30.

38. Vacca P, Vitale C, Munari E, Cassatella MA, Mingari MC, Moretta L. Human Innate Lymphoid Cells: Their Functional and Cellular Interactions in Decidua. Front Immunol 2018; 9: 1897.

39. Oksana Shynlova AB-R, Tali Farine, Kristina M. Adams Waldorf, Caroline Dunk and Stephen J. Lye. Decidual Inflammation Drives Chemokine-Mediated Immune Infiltration Contributing to Term Labor. J Immunol 2021; 207(8): 2015–26.

40. Efremova M, Vento-Tormo M, Teichmann SA, Vento-Tormo R. CellPhoneDB: inferring cell-cell communication from combined expression of multi-subunit ligand-receptor complexes. Nat Protoc 2020; 15(4): 1484–506.

41. Shuya LL, Menkhorst EM, Yap J, Li P, Lane N, Dimitriadis E. Leukemia inhibitory factor enhances endometrial stromal cell decidualization in humans and mice. PLoS One 2011; 6(9): e25288.

42. Wiesolek HL, Bui TM, Lee JJ, et al. Intercellular Adhesion Molecule 1 Functions as an Efferocytosis Receptor in Inflammatory Macrophages. Am J Pathol 2020; 190(4): 874–85.

43. Kramer AC, Erikson DW, McLendon BA, et al. SPP1 expression in the mouse uterus and placenta: implications for implantationdagger. Biol Reprod 2021; 105(4): 892–904.

44. Gomes AO, Barbosa BF, Franco PS, et al. Macrophage Migration Inhibitory Factor (MIF) Prevents Maternal Death, but Contributes to Poor Fetal Outcome During Congenital Toxoplasmosis. Front Microbiol 2018; 9: 906.

45. Devergne O, Coulomb-L’Hermine A, Capel F, Moussa M, Capron F. Expression of Epstein-Barr virus-induced gene 3, an interleukin-12 p40-related molecule, throughout human pregnancy: involvement of syncytiotrophoblasts and extravillous trophoblasts. Am J Pathol 2001; 159(5): 1763–76.

46. Cheng JC, Chang HM, Leung PC. Transforming growth factor-beta1 inhibits trophoblast cell invasion by inducing Snail-mediated down-regulation of vascular endothelial-cadherin protein. J Biol Chem 2013; 288(46): 33181–92.

47. Rouillard AD, Gundersen GW, Fernandez NF, et al. The harmonizome: a collection of processed datasets gathered to serve and mine knowledge about genes and proteins. Database (Oxford) 2016; 2016.

48. Poli-Neto OB, Meola J, Rosa ESJC, Tiezzi D. Transcriptome meta-analysis reveals differences of immune profile between eutopic endometrium from stage I-II and III-IV endometriosis independently of hormonal milieu. Sci Rep 2020; 10(1): 313.

49. Petracco R, Grechukhina O, Popkhadze S, Massasa E, Zhou Y, Taylor HS. MicroRNA 135 regulates HOXA10 expression in endometriosis. J Clin Endocrinol Metab 2011; 96(12): E1925–33.

50. Matalliotakis M, Zervou MI, Matalliotaki C, et al. The role of gene polymorphisms in endometriosis. Mol Med Rep 2017; 16(5): 5881–6.

51. Martinez-Fierro ML, Hernadez-Delgadillo GP, Flores-Mendoza JF, et al. Fibroblast Growth Factor Type 2 (FGF2) Administration Attenuated the Clinical Manifestations of Preeclampsia in a Murine Model Induced by L-NAME. Front Pharmacol 2021; 12: 663044.

52. Mrozikiewicz AE, Kurzawinska G, Gozdziewicz-Szpera A, et al. Effects of TIMP1 rs4898 Gene Polymorphism on Early-Onset Preeclampsia Development and Placenta Weight. Diagnostics (Basel) 2022; 12(7).

53. Yang D, Dai F, Yuan M, et al. Role of Transforming Growth Factor-beta1 in Regulating Fetal-Maternal Immune Tolerance in Normal and Pathological Pregnancy. Front Immunol 2021; 12: 689181.

54. Brooks SA, Martin E, Smeester L, Grace MR, Boggess K, Fry RC. miRNAs as common regulators of the transforming growth factor (TGF)-beta pathway in the preeclamptic placenta and cadmium-treated trophoblasts: Links between the environment, the epigenome and preeclampsia. Food Chem Toxicol 2016; 98(Pt A): 50–7.

55. Moufarrej MN, Vorperian SK, Wong RJ, et al. Early prediction of preeclampsia in pregnancy with cell-free RNA. Nature 2022; 602(7898): 689-94.

56. Keskin DB, Allan DS, Rybalov B, et al. TGFbeta promotes conversion of CD16+ peripheral blood NK cells into CD16- NK cells with similarities to decidual NK cells. Proc Natl Acad Sci U S A 2007; 104(9): 3378–83.

57. Cerdeira AS, Rajakumar A, Royle CM, et al. Conversion of peripheral blood NK cells to a decidual NK-like phenotype by a cocktail of defined factors. J Immunol 2013; 190(8): 3939–48.

58. Burton GJ. Oxygen, the Janus gas; its effects on human placental development and function. J Anat 2009; 215(1): 27–35.

59. Burton GJ, Cindrova-Davies T, Yung HW, Jauniaux E. HYPOXIA AND REPRODUCTIVE HEALTH: Oxygen and development of the human placenta. Reproduction 2021; 161(1): F53–F65.

60. Moffett A, Shreeve N. Local immune recognition of trophoblast in early human pregnancy: controversies and questions. Nat Rev Immunol 2022: 1–14.

61. Pollheimer J, Vondra S, Baltayeva J, Beristain AG, Knofler M. Regulation of Placental Extravillous Trophoblasts by the Maternal Uterine Environment. Front Immunol 2018; 9: 2597.

62. Benny PA, Alakwaa FM, Schlueter RJ, Lassiter CB, Garmire LX. A review of omics approaches to study preeclampsia. Placenta 2020; 92: 17–27.

63. Tarca AL, Romero R, Benshalom-Tirosh N, et al. The prediction of early preeclampsia: Results from a longitudinal proteomics study. PLoS One 2019; 14(6): e0217273.

64. MacDonald TM, Walker SP, Hannan NJ, Tong S, Kaitu’u-Lino TJ. Clinical tools and biomarkers to predict preeclampsia. EBioMedicine 2022; 75: 103780.

65. Rodriguez-Rivera C, Garcia MM, Molina-Alvarez M, Gonzalez-Martin C, Goicoechea C. Clusterin: Always protecting. Synthesis, function and potential issues. Biomed Pharmacother 2021; 134: 111174.

66. Jia T, Jacquet T, Dalonneau F, et al. FGF-2 promotes angiogenesis through a SRSF1/SRSF3/SRPK1-dependent axis that controls VEGFR1 splicing in endothelial cells. BMC Biol 2021; 19(1): 173.

67. Tang J, Kang Y, Huang L, Wu L, Peng Y. TIMP1 preserves the blood-brain barrier through interacting with CD63/integrin beta 1 complex and regulating downstream FAK/RhoA signaling. Acta Pharm Sin B 2020; 10(6): 987–1003.

68. Kim HJ, Yoo EK, Kim JY, et al. Protective role of clusterin/apolipoprotein J against neointimal hyperplasia via antiproliferative effect on vascular smooth muscle cells and cytoprotective effect on endothelial cells. Arterioscler Thromb Vasc Biol 2009; 29(10): 1558–64.

69. Lebrin F, Deckers M, Bertolino P, Ten Dijke P. TGF-beta receptor function in the endothelium. Cardiovasc Res 2005; 65(3): 599–608.

